# Rabies anterograde monosynaptic tracing reveals organization of spinal sensory circuits

**DOI:** 10.1101/2021.02.11.429920

**Authors:** Sofia Pimpinella, Niccolò Zampieri

## Abstract

Somatosensory neurons detect vital information about the environment and internal status of the body, such as temperature, touch, itch and proprioception. The circuit mechanisms controlling the coding of somatosensory information and the generation of appropriate behavioral responses are not clear yet. In order to address this issue, it is important to define the precise connectivity patterns between primary sensory afferents dedicated to the detection of different stimuli and recipient neurons in the central nervous system. In this study we used a rabies tracing approach for mapping spinal circuits receiving sensory input from distinct, genetically defined, modalities. We analyzed the anatomical organization of spinal circuits involved in coding of thermal and mechanical stimuli and showed that somatosensory information from distinct modalities is relayed to partially overlapping ensembles of interneurons displaying stereotyped laminar organization, thus highlighting the importance of positional features and population coding for the processing and integration of somatosensory information.

## Introduction

The somatosensory system is responsible for detecting a wide variety of sensory information and generate appropriate behavioral responses. The circuit mechanisms controlling the detection of different modalities and its transformation into motor actions are not completely understood. Much progress has been made in the characterization of primary somatosensory neurons in the peripheral nervous system, and physiological and molecular descriptions of different subtypes specialized in the detection of discrete modalities exist (Abraira and Ginty, 2013; Vriens et al., 2014; Le Pichon and Chesler, 2014; Zampieri and De Nooij, 2020). However, comparatively little is known about the logic underlying the coding of sensory information and the generation of appropriate motor behaviors.

The specialization of peripheral afferents for the detection of distinct stimuli represents the foundation underlying the specificity theory, which proposes that different sensory information is encoded along parallel dedicated pathways or labelled lines (Norrsell et al, 1999). An alternative view, supported by studies on pain, is based on pattern theory and it postulates that perception is generated by temporal summation of various peripheral inputs at the level of relay centers in the central nervous system (CNS; Perl, 2007). More recently, a synergistic model, population coding, has been proposed (Ma, 2010). It suggests that cross-talk between labelled lines in the CNS is responsible for coding of sensory perception. This hypothesis highlights the functional specialization of primary sensory afferents and postulate the existence of specific patterns of connectivity with second order neurons in the spinal cord and medulla. Thus, defining the location and identity of spinal interneurons receiving input from distinct sensory modalities represent an important step toward resolving the circuit mechanisms controlling the coding of somatic sensation. However, systematic analysis has been so far precluded by the lack of high-throughput approaches that directly links the subtype identity of primary afferents with their spinal targets (Bokiniec et al., 2018).

Transsynaptic tracing using rabies virus is a powerful tool for mapping neural circuits in the brain and spinal cord (Wickersham et al., 2007; Callaway and Luo, 2015). Rabies virus has also been shown to infect primary sensory neurons in the peripheral nervous system and move in the anterograde direction to spread into synaptic targets into the CNS (Ugolini, 2010; Velandia-Romero et al., 2013; Bauer et al., 2014). However, several limitations have hampered the use of rabies tracing for systematic analysis of spinal sensory circuits. First, not all sensory afferents are susceptible to rabies virus infection (Albisetti et al., 2017). Second, neuronal infection after peripheral, intramuscular or cutaneous, rabies injection is only efficient in neonatal mice (Stepien et al., 2010; Zampieri et al., 2014; Zhang et al., 2015). Finally, questions have been raised about the directionality of the transfer and whether it represents bona fide anterograde tracing of postsynaptic targets or retrograde labeling of presynaptic neurons providing axo-axonic input to sensory afferents (Zhang et al., 2015).

In this study, we show that intraspinal injection of EnvA-pseudotyped rabies virus can retrogradely infect TVA expressing neurons in the dorsal root ganglia (DRG) without any obvious limitation in either tropism or timing, and jump in the anterograde direction into second order spinal neurons allowing high resolution mapping of post-sensory circuits. Analysis of proprioceptive circuits resulted in identification of motor neurons and interneuron subtypes, that are well known post-synaptic partners, thus indicating genuine anterograde transsynaptic transfer. We applied this approach to identify spinal targets of sensory afferents detecting thermal and mechanical stimuli and observed a high degree of spatial segregation along dorso-ventral axis of the spinal cord. However, we identified convergence of distinct post-sensory circuits in mutual exclusive areas in dorsal laminae I-III, thus indicating important roles as hubs for integration of multiple sensory modalities. These findings highlight the functional relevance of the laminar organization of the spinal cord and emphasize the essential role of positional features, as a determinant in the assembly and function of sensory-motor circuits.

## Results

### Retrograde infection of primary somatosensory neurons and anterograde transfer into second order neurons

In order to characterize the anatomical organization of spinal circuits according to the somatosensory input they receive, we combined mouse genetic and rabies virus (RV, SAD B19 strain; Wickersham et al., 2007) monosynaptic tracing techniques (Figure 1A). To achieve cellular specificity in RV infection and subsequent monosynaptic transfer we used a mouse line driving conditional expression of the TVA receptor, the RV glycoprotein (G) and a nuclear GFP reporter in combination with Cre lines targeting defined subsets of somatosensory neurons (*Rosa26^Lox-stop-LoxHTB^* or *HTB*; Li et al., 2013). Expression of the TVA receptor and G are required for selective infection by EnvA pseudotyped G-deleted RV (RVΔG-mCherry/EnvA) and subsequent monosynaptic spreading, while the nuclear GFP reporter allows identification of starter cells (Figure 1A). We first focused on proprioceptive circuits that have been quite extensively characterized at anatomical and physiological levels (Zampieri et al., 2014; Balaskas et al., 2020). Thus, we crossed *parvalbumin::Cre* (*PV::Cre*), which is expressed in proprioceptive sensory neurons and a subset of low-threshold mechanoreceptors (LTMR) with the *HTB* mouse line (Hippenmeyer et al., 2007; de Nooij et al., 2013). First, we confirmed the expression specificity of the *HTB* allele and found that at lumbar levels about 96% of GFP^+^ DRG neurons were also PV^+^ (Figure 1B and C). In addition, GFP was not detected in the lumbar spinal cord up to postnatal (p) day 10, indicating that neither the TVA receptor nor G are expressed in spinal neurons up to this stage (Figure S1A). In order to target sensory neurons independently of their subtype identity and peripheral target connectivity we delivered RV directly in the spinal cord to gain access to somatosensory afferents (Figure 1A). Unilateral stereotactic injection of RVΔG-mCherry/EnvA at lumbar (L) level 1 of p9 *PV::Cre^+/−^; HTB^f/f^* (hereafter referred to as *PV^HTB^*) mice resulted in the infection of PV^+^ neurons in L2-L4 DRG (Figures 1A, D, E, F, and S1B). Next, we examined the spinal cord seven days after rabies injection and observed labeling of interneurons and motor neurons (Figure 1G and Video S1). In contrast, when we injected *PV::Cre^+/−^; HTB^f/+^* mice, we obtained infection of PV^+^ DRG neurons but negligible labeling of spinal neurons (Figure S1C). This observation indicates that one copy of the *HTB* allele can promote sufficient expression of TVA to drive interaction with EnvA pseudotyped RV but not enough G to support transsynaptic transfer. Strikingly, we observed that the majority of neurons labeled in the spinal cord were located along or nearby the axonal trajectories of proprioceptive sensory afferents (Figures 1G and 1H). Altogether these data indicate that spinal injection of rabies results in retrograde infection of proprioceptive neurons and anterograde transsynaptic spreading into neuronal targets in the spinal cord.

**Figure 1.**
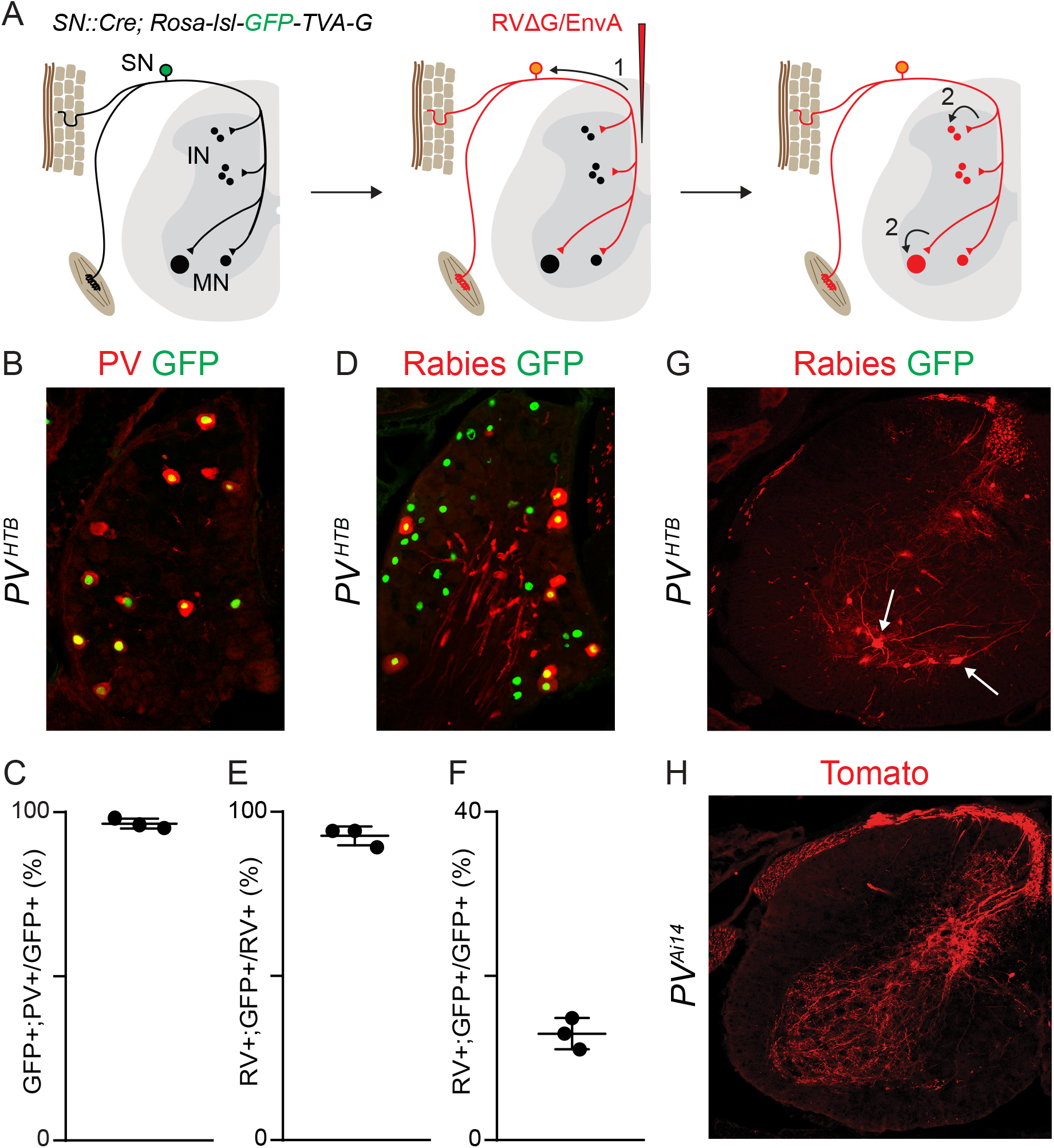
Retrograde infection of primary somatosensory neurons and anterograde monosyaptic spread into spinal neurons. A) Schematics representing the strategy for genetic targeting of G and TVA expression in DRG neurons and monosynaptic tracing with pseudotyped rabies injection in the spinal cord. SN, sensory neurons; IN, interneurons; MN, motor neurons; 1, primary infection; 2, secondary infection. B) Parvalbumin expression in GFP^+^ sensory neurons in p9 *PV^HTB^* mice. C) Specificity of genetic tracing with the *PV^HTB^* line expressed as a percentage of GFP^+^ sensory neurons labeled by parvalbumin staining. D) Rabies expression (mCherry) in GFP^+^ sensory neurons after RVΔG-mCherry/EnvA injection in *PV^HTB^* mice. E) Specificity of sensory neurons targeting expressed as a percentage of RV^+^ sensory neurons labeled by nuclear GFP after RVΔG-mCherry/EnvA injection in *PV^HTB^* mice. F) Efficiency of sensory neurons targeting expressed as a percentage of GFP^+^ sensory neurons labeled by mCherry after RVΔG-mCherry/EnvA injection in *PV^HTB^* mice. G) Rabies expression (mCherry) in spinal neurons at p16 after RVΔG-mCherry/EnvA injection in p9 *PV^HTB^* mice. Arrows points to motor neurons in the ventral spinal cord. H) tdTomato labeling of proprioceptive sensory afferents in the spinal cord of *PV::Cre^+/−^; Ai14^f/+^*mice.

### Organization and identity of second order neurons receiving proprioceptive input

We next sought to characterize the spinal neurons labeled by rabies tracing (Figure 2A). We generated three-dimensional maps of infected neurons and analyzed their positional organization in the spinal cord. The vast majority of second order neurons were located ipsilateral to the point of injection (Figures 2B, 2C and S2). Distribution along the dorso-ventral axis presented three distinct peaks corresponding to the dorsal, intermediate and ventral spinal cord (Figures 2B and 2D). Consistent with unbiased access to sensory afferents independent of their peripheral target, we found rabies-labelled motor neurons of both lateral and medial motor column identity in the ventral ipsilateral side (Figures 2A, 2C and S2). The connectivity patterns obtained from different tracing experiments were qualitatively and quantitatively reproducible as shown in individual maps, distribution and correlation analyses (“IN vs IN” r ≥ 0.9; “MN vs MN” r ≥ 0.8; Figures 2D, 2H and S2). We observed some variability in the amount of neuronal labelling in different experiments, however the ratio between the number of starter cells and second order neurons, defined as the “connectivity index”, remained constant indicating that, under these conditions, rabies can reproducibly label ~5 spinal neurons for each primary sensory neuron infected (Figures 2E-G and Table S1). Interestingly, a similar level of transsynaptic transfer was previously reported in retrograde tracing experiments from motor neurons using the SAD B19 strain (Reardon et al., 2016). Next, we investigated the identity of spinal neurons labelled by rabies virus. Aside from motor neurons, several cardinal classes of spinal neurons are known to receive direct proprioceptive input (Eccles et al., 1957; Côté et al., 2018). We analysed the expression of markers that, along with positional information, define V2a, V1, V0 and dI4 identities at early postnatal stages (Zampieri et al., 2014; Bikoff et al., 2016). We found rabies labelled Chx10^+^ V2a interneurons, FoxP2^+^ V1 interneurons, ventrally positioned calbindin^+^ Renshaw cells, and Lhx1^+^interneurons whose dorsal position is suggestive of dI4 identity (Figure S3). These findings confirm that rabies labels spinal neurons that are known to receive monosynaptic proprioceptive input and therefore represent genuine postsynaptic targets.

**Figure 2.**
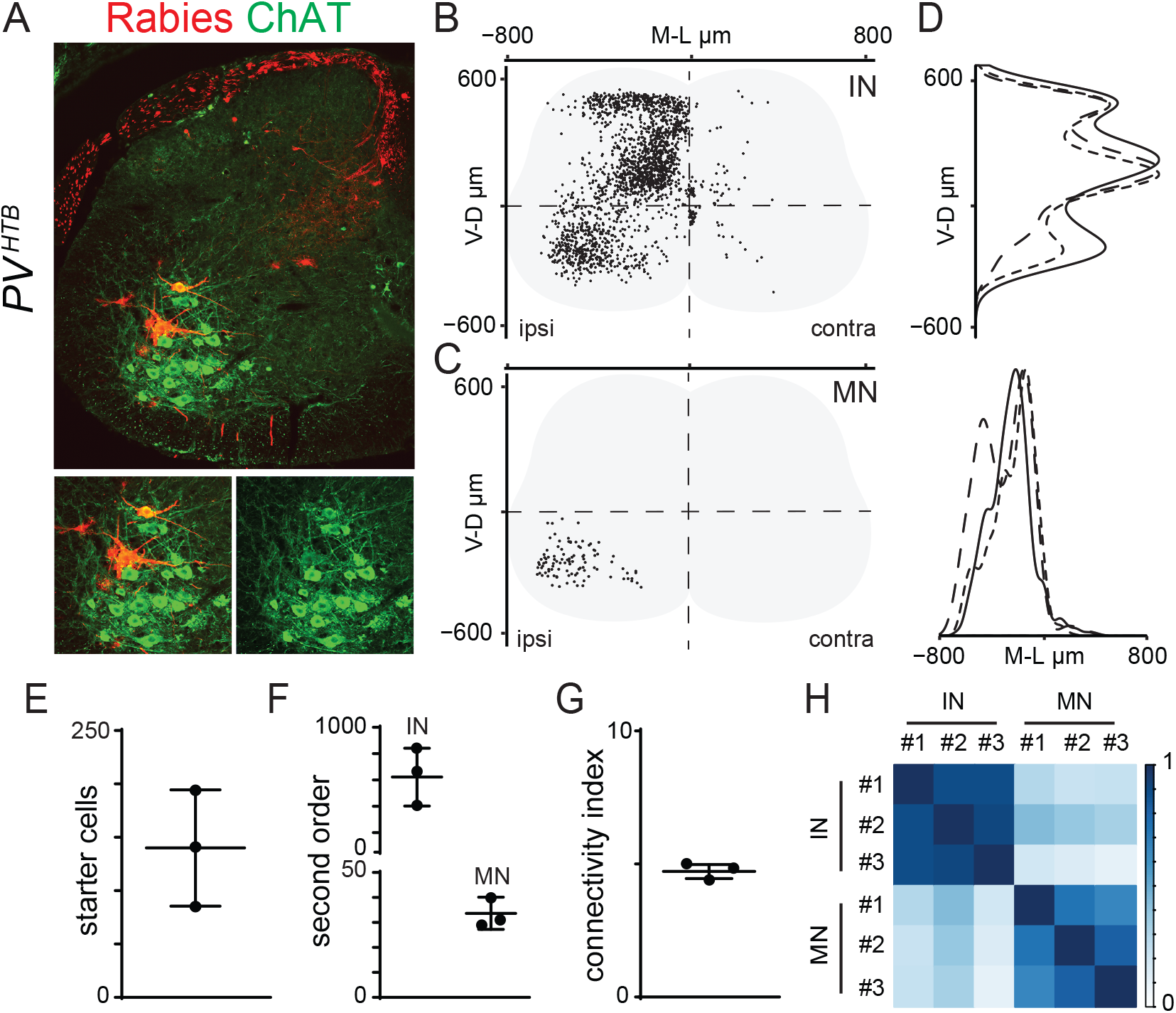
Rabies tracing of spinal proprioceptive circuits. A) Rabies expression (mCherry) in spinal interneurons and ChAT^+^ motor neurons after RVΔG-mCherry/EnvA injection in p16 *PV^HTB^* mice. B) Digital reconstruction of RV^+^ interneuron positions in *PV^HTB^* experiments. IN, interneurons. C) Digital reconstruction of RV^+^; ChAT^+^ motor neuron positions in *PV^HTB^* experiments. MN, motor neurons. D) Dorso-ventral (top) and medio-lateral (bottom) density analyses of RV^+^ interneurons (top) and RV^+^; ChAT^+^ motor neurons (bottom) in three *PV^HTB^* experiments. E) Number of starter cells defined as GFP^+^; RV^+^ sensory neurons in *PV^HTB^* experiments. F) Number of spinal neurons traced in *PV^HTB^* experiments. IN, interneurons; MN, motor neurons. G) Connectivity index, the average number of second order neurons traced from a single starter cell in *PV^HTB^* experiments. H) Correlation analysis of interneurons and motor neurons positional coordinates in *PV^HTB^* experiments (“IN vs IN” R≥ 0.9; “MN vs MN” R≥ 0.8). Scale bar indicates correlation values. IN, interneurons; MN, motor neurons.

### Anterograde tracing from thermosensitive neurons

In order to explore the anatomical organization of spinal somatosensory circuits we mapped post-sensory neurons receiving input from different primary afferents. Since the *PV::Cre* line gives access to neurons of mechanoreceptive lineage, mainly proprioceptors and a subset of LTMR, we decided to focus on thermosensation, a distinct modality that is accessible using available mouse genetic tools. We took advantage of the *TRPV1::Cre* and *TRPM8::Cre* mouse lines that are known to target hot- and cold-sensing DRG neurons (Mishra et al., 2011; Yarmolinsky et al., 2016). *TRPV1* is transiently expressed during development by most sensory neurons of thermoceptive lineage, while *TRPM8* is restricted to a smaller subset of cold-sensing neurons (Dhaka et al., 2008; Mishra et al., 2011). Indeed, GFP expression in *TRPV1::Cre^+/−^; HTB^f/f^* and *TRPM8::Cre^+/−^; HTB^f/f^* (hereafter referred to as *TRPV1^HTB^* and *TRPM8^HTB^*) mice revealed a clear difference in the labeling of DRG neurons (Figures 3A and B). This observation was confirmed by tracing sensory afferents, in *TRPV1::Cre^+/−^; Ai14^f/+^* mice we found dense staining in the dorsal spinal cord while only sparse signal was detected in the case of *TRPM8::Cre^+/−^; Ai14^f/+^* (Figures S4C-D). In both cases we did not detect any labeling of spinal neurons with either the *HTB* or the *Ai14* reporter lines(Figures 3E, 3F and S4C-D). As previously done for *PV^HTB^* experiments, we performed L1 unilateral stereotactic injection of RVΔG-mCherry/EnvA in p9 mice and performed analysis at p16. In both cases we obtained selective infection of GFP^+^ DRG neurons (Figures 3A-C). We observed similar efficiencies in primary infection, however, because of the difference in the amount of sensory neurons expressing Cre in the *TRPV1::Cre* and *TRPM8::Cre* lines, the number of starter cells was much higher in *TRPV1^HTB^* experiments (Figure 3A, 3B, 3D, 3I and Table S1). Surprisingly, we did not observe a proportional increase in the number of second order neurons labeled in *TRPV1^HTB^* mice, thus resulting in a low connectivity index (Figures 3I-K and Table S1). Next, we examined the spinal cords of rabies injected *TRPV1^HTB^* and *TRPM8^HTB^* mice and found extensive labeling in the ipsilateral side with higher incidence of RV^+^ neurons in the dorsal horn that sharply decreased in the intermediate and ventral spinal cord (Figures 3E-H). The overall spatial organization of RV^+^ neurons in *TRPV1^HTB^* and *TRPM8^HTB^* experiments were qualitatively similar and injections reproducible, as shown by single maps, distribution, and correlation analyses (Figures S4A-B, S4E). These data show that rabies can be used to trace from distinct primary somatosensory neuron subtypes and indicate that spinal sensory circuits encoding for different modalities, such as thermosensation and proprioception, are kept mostly separated, highlighting the functional specialization of the dorsal and ventral spinal cord in the control of sensory processing and motor control.

**Figure 3.**
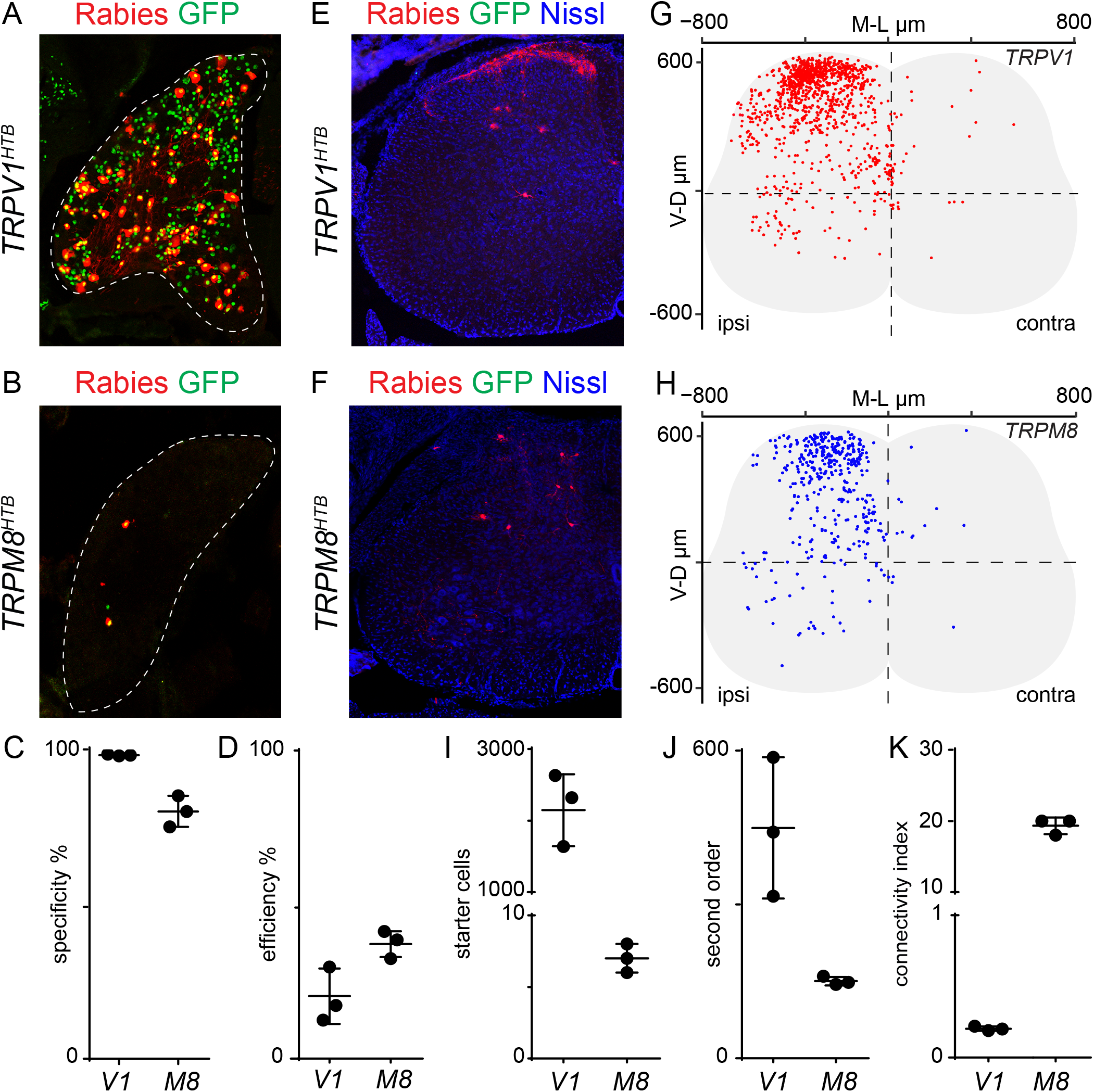
Rabies tracing of spinal thermosensitive circuits. A and B) Rabies expression (mCherry) in GFP^+^ sensory neurons labeled after RVΔG-mCherry/EnvA injection in *TRPV1^HTB^* (A) and *TRPM8^HTB^* (B) mice. C) Specificity of sensory neurons targeting expressed as a percentage of RV^+^ sensory neurons labeled by nuclear GFP after RVΔG-mCherry/EnvA injection in *TRPV1^HTB^* and *TRPM8^HTB^* mice. D) Efficiency of sensory neurons targeting expressed as a percentage of GFP^+^ sensory neurons labeled by mCherry after RVΔG-mCherry/EnvA injection in *TRPV1^HTB^* and *TRPM8^HTB^* mice. E and F) RV^+^ (mCherry) spinal neurons after RVΔG-mCherry/EnvA injection in *TRPV1^HTB^* (E) and *TRPM8^HTB^* (F) mice. G and H) Digital reconstruction of RV^+^ interneuron positions in *TRPV1^HTB^* (G) and *TRPM8^HTB^* (H) experiments. I) Number of starter cells defined as GFP^+^; RV^+^ sensory neurons in *TRPV1^HTB^* and *TRPM8^HTB^* experiments. J) Number of spinal neurons traced in *TRPV1^HTB^* and *TRPM8^HTB^* experiments. K) Connectivity index defined as the average number of second order neurons traced from a single starter cell in *TRPV1^HTB^* and *TRPM8^HTB^* experiments.

### Organization of post-sensory circuits in the dorsal laminae of the spinal cord

Recent studies demonstrated the importance of topographic organization of dorsal spinal interneurons for encoding reflexes mediated by inflammatory and noxious stimuli (Gatto et al., 2021; Peirs et al., 2021). Thus, we asked whether the distribution of neurons labelled in *PV^HTB^*, *TRPV1^HTB^,* and *TRPM8^HTB^* may reveal the anatomical basis for the functional specificity of spinal somatosensory circuits. In *PV^HTB^* experiments we found 43% of rabies-labeled neurons in the intermediate spinal cord (defined as the dorso-ventral area from 0 to 300μm) and a similar number of cells in the ventral (23%; 0 to −600μm) and dorsal (29%; 300 to 600μm) areas (Figure 4A). In contrast, the majority of neurons traced after rabies injections in *TRPV1^HTB^* and *TRPM8^HTB^* mice were located in the dorsal spinal cord (Figure 4A; *TRPV1^HTB^* = 78% and *TRPM8^HTB^* = 65%). Correlation analysis confirmed this observation by showing that Cartesian coordinates of RV^+^ neurons in *TRPV1^HTB^* and *TRPM8^HTB^* experiments highly correlate with each other but not with the ones from *PV^HTB^* (“*TRPV1^HTB^* vs *TRPM8^HTB^*” r ≥ 0.85; “*TRPV1^HTB^*or *TRPM8^HTB^* vs *PV^HTB^*” r ≤ 0.55; Figure 4B). Despite the broad dorso-ventral segregation in the distributions of neurons receiving thermal and mechanical information, an area of potential overlap is evident in the dorsal spinal cord (Figure 4A). We used staining for PKCγ, a marker for lamina IIi and III (Polgar et al., 1999), as an internal reference for assessing relative positioning of dorsal interneurons labelled in *PV^HTB^*, *TRPV1^HTB^,* and *TRPM8^HTB^* experiments (Figure 4C). The data indicate that in *PV^HTB^* experiments RV^+^ neurons are rarely found above lamina IIi, as opposed to *TRPV1^HTB^* mice where RV^+^ neurons are located mostly in lamina I and IIo, largely overlapping with the CRGP termination zone (Figures 4C-E and G). In addition, in *PV^HTB^* experiments we observed a subset of neurons displaying prominent laminar positioning mostly overlapping with PKCγ labeling, an area known to receive extensive input from cutaneous LTMR (Figures 4C, 4D and 4E; Abraira et al., 2017). In contrast, spinal neurons traced in *TRPM8^HTB^* mice presented a more homogenous distribution across dorsal layers, thus resulting in one area of partial overlap with *TRPV1^HTB^* traced neurons in laminae I-IIo and one of partial overlap with *PV^HTB^* traced neurons in laminae IIi-III (Figure 4F). Altogether, these data indicate that interneurons residing in the superficial laminae can be divided into at least three different populations according to the sensory input they receive.

**Figure 4.**
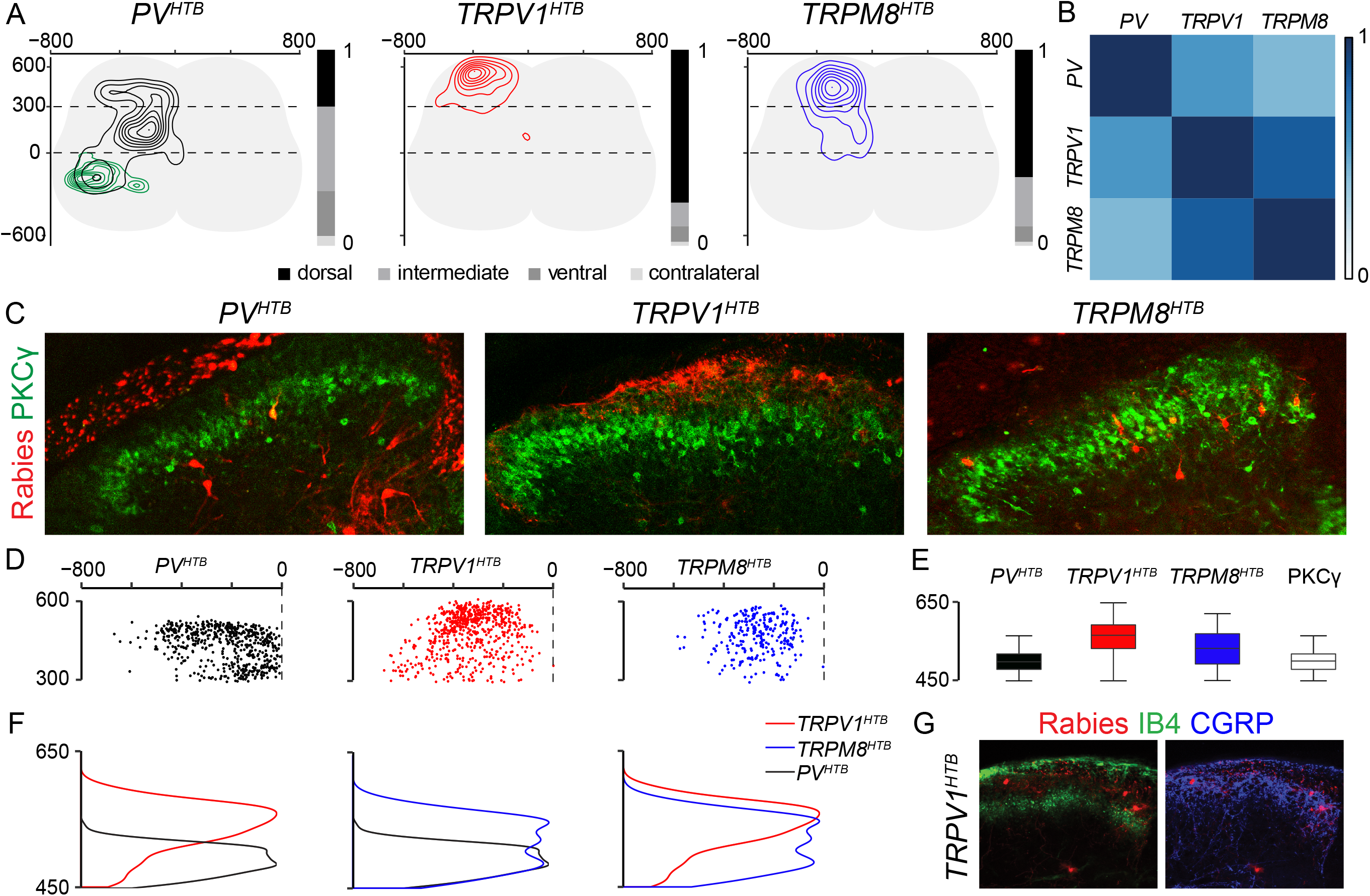
Organization of sensory circuits for mechanical and thermal sensation in the spinal cord. A) Transverse contour density plots and dorso-ventral distribution of post-sensory neurons in the dorsal (300 to 600μm), intermediate (0 to 300μm), ventral (0 to −600μm) and contralateral spinal cord of *PV^HTB^* (interneurons: black; motor neurons: green)*, TRPV1^HTB^* (Red) and *TRPM8^HTB^* (Blue) experiments. B) Correlation analysis of post-sensory neurons Cartesian coordinates in *PV^HTB^, TRPV1^HTB^* and *TRPM8^HTB^* experiments (“*TRPV1^HTB^* vs *TRPM8^HTB^*” r ≥ 0.85; “*TRPV1^HTB^*or *TRPM8^HTB^* vs *PV^HTB^*” r ≤ 0.55). Scale bar indicates correlation values. C) PKCγ and mCherry expression in ipsilateral dorsal spinal cords after RVΔG-mCherry/EnvA injection in *PV^HTB^, TRPV1^HTB^* and *TRPM8^HTB^* mice. D) Digital reconstruction of RV^+^ interneuron positions in the dorsal spinal cord of *PV^HTB^* (black), *TRPV1^HTB^* (red) and *TRPM8^HTB^* experiments. E) Box-plot showing dorso-ventral the distributions of RV^+^ interneurons in the dorsal horn of *PV^HTB^* (black), *TRPV1^HTB^* (red) and *TRPM8^HTB^* (blue) experiments. PKCγ staining (white) is used as an internal reference. F) Dorso-ventral density plots showing the distributions of RV^+^ interneurons in the dorsal horn of *PV^HTB^* (black), *TRPV1^HTB^* (red) and *TRPM8^HTB^* (blue) experiments. G) CGRP (laminae I and IIo), IB4 (lamina IIm), and mCherry expression in ipsilateral dorsal spinal cords after RVΔG-mCherry/EnvA L1 injection in p9 *TRPV1^HTB^* mice.

## Discussion

In order to understand the functional organization of spinal circuits controlling the processing of sensory information, it is critical to determine the patterns of connectivity between distinct primary sensory neuron subtypes and their targets in the central nervous system. In this study, by combining mouse genetics and rabies monosynaptic tracing techniques we identified an approach to directly link sensory input from defined, modality specific, primary afferents to neuronal targets in the spinal cord and analyzed the anatomical organization of spinal circuits encoding thermal and mechanical information.

The approach takes advantage of the ability of primary sensory neurons to support rabies transsynaptic transfer in the anterograde direction (Ugolini, 2010; Velandia-Romero et al., 2013; Bauer et al., 2014; Zampieri et al., 2014). In contrast to previous studies that used peripheral rabies injection, either cutaneous or intramuscular, to infect sensory neurons through their terminals, we opted for stereotactic injection of EnvA pseudotyped rabies virus in the spinal cord to infect TVA-expressing neurons through their central afferents. This route has two main advantages. First, the efficiency of rabies infection of DRG neurons, via their peripheral terminals is known to decrease rapidly within the first neonatal days, essentially restricting the experimental window from p1-p4 (Zampieri et al., 2014; Zhang et al., 2015). This limitation does not apply to intraspinal injection, thus opening the way for studying the organization of spinal post-sensory circuits during and after postnatal development in physiological or disease models. Second, intraspinal injection allows unbiased access to all sensory afferents projecting at a desired spinal level independent of their identity or pattern of peripheral innervation (i.e.: hairy vs glabrous skin, cutaneous vs muscle, etc.), in principle allowing direct comparisons of post-sensory circuits from different modalities without any limitation.

We used a mouse genetic strategy to specify starter cells by driving conditional expression of the TVA receptor, G protein and a reporter under control of Cre recombinase. This is an effective and relatively simple method for driving transgene expression in all neurons of interest. However, it requires a high degree of specificity in the Cre line, otherwise transient or leaky expression could result in the generation of multiple sets of cells capable of supporting rabies infection and transsynaptic tracing. For this reason, we carefully analyzed the patterns of the nuclear GFP and tdTomato reporters in the DRG and spinal cord of the Cre lines employed in this study. In order to ensure more stringent specificity, intersectional genetic and viral strategies can be used. For example, complementation of TVA and G expression using peripheral AAV injection in combination with rabies intraspinal delivery could eliminate specificity issues common to many Cre lines.

In agreement with a previous report, we did not find any obvious restriction in the ability of EnvA-pseudotyped rabies virus to infect TVA expressing somatosensory neuron (Albisetti et al., 2017). However, in comparison to TRPM8 and PV experiments, we observed relatively low connectivity when tracing from *TRPV1^HTB^* mice. Our data do not allow to distinguish whether this observation reflects an intrinsic property of these circuits or could hint at a limited ability of a subset of *TRPV1::Cre* neurons to support rabies spreading. It has been suggested that neural activity may have an important role in promoting efficient rabies transsynaptic transfer. Many nociceptors are labelled by the *TRPV1::Cre* line that because of the controlled conditions of laboratory mouse housing may not be active, thus possibly limiting their contributions to rabies tracing. A similar scarcity in connectivity has been previously shown in tracing experiments from *TRPV1::Cre* sensory neurons after rabies cutaneous injection (Zhang et al., 2015). The authors interpreted their results as an indication that transsynaptic labeling from sensory neurons represents retrograde transfer into presynaptic neurons through relatively infrequent axo-axonic synapses instead of anterograde transfer into postsynaptic targets (Zhang et al., 2015). Analysis of neuronal identity and position in *PV^HTB^* experiments strongly support anterograde transfer into postsynaptic targets, as we consistently observe labeling of motor neurons and spinal interneurons that are well-known recipients of monosynaptic input from proprioceptive sensory afferents (Eccles et al., 1957; Zampieri et al., 2014; Bikoff et al., 2016; Côté et al., 2018).

In order to reveal the anatomical organization of spinal somatosensory circuits, we used three different mouse lines, *PV::Cre* to label proprioceptive neurons and a subset of LTMR, *TRPV1::Cre* to label thermosensitive neurons, and *TRPM8::Cre* to label cold-sensing neurons (Hippenmeyer et al., 2007; Mishra et al., 2011; Yarmolinsky et al., 2016). We were therefore able to map neurons involved in the detection of two different stimulus modalities, proprioception and thermosensation, as well as circuits for more defined sensory features, cutaneous mechanoreceptors and cold sensing neurons. Positional analysis of post-sensory neurons revealed shared and distinct features of spinal somatosensory circuits. First, in all cases analyzed, we observed a very prominent ipsilateral bias in connectivity, with very limited labeling of contralateral neurons, indicating that the first relay stations processing somatic sensation do not directly integrate information coming from both sides of the body. Second, post-sensory neurons receiving thermal and proprioceptive information are mostly segregated along the dorso-ventral axis highlighting the functional separation of the dorsal and ventral spinal cord for sensory processing and motor control, respectively. Third, at a finer level of resolution, the anatomical organization of post-sensory circuits in the dorsal horn reflects the recently described functional specialization of superficial spinal interneurons in laminae I-IIo for encoding reflexes mediated by inflammatory and noxious stimuli, and of deeper interneurons in laminae IIi-IV for sensory-motor behaviours driven by mechanical inputs (Gatto et al., 2021; Peirs et al., 2021). Interneurons labeled in *TRPV1^HTB^* experiments, that include afferents detecting noxious thermal stimuli are present at higher density in lamina I and IIo, neatly segregated from the ones traced in *PV^HTB^* experiments, representing inputs relaying proprioceptive and cutaneous mechanoreceptive information, that are found in deeper layers starting from lamina IIi. Interestingly, spinal targets of afferents traced in *TRPM8^HTB^* experiments, that detect non noxious thermal information, present a more homogenous distribution throughout the dorsal horn, selectively overlapping with TRPV1-labelled neurons in laminae I-IIi and PV output areas in laminae IIo-III, thus indicating that even somatosensory information coming from distinct modalities is not strictly kept separated at the level of first-order spinal neurons. Altogether, these findings support a population coding model where different, modality specific, sensory inputs converge on ensembles of spinal interneurons that present stereotyped spatial organization and control different sensory-motor functions (Gradwell and Abraira, 2021).

## Acknowledgements

We would like to thank Mark Hoon (NIH, USA) for generously providing the *TRPV1::Cre* and *TRPM8::Cre* mouse lines; Martyn Goulding (Salk Institute, USA) for the *Rosa-lsl-HTB* mouse line. Carmen Birchmeier for generously sharing the anti Lbx1 and anti CGRP antibodies, Susan Morton for the anti Lhx1 antibody. Stephan Dietrich for helping with tissue clearing and light sheet microscopy. Liana Kosizki for technical assistance and the MDC Advanced Light Microscope facility for assistance with image acquisition and analysis. We are grateful to Marco Beato, Jay Bikoff, Joriene de Nooij, Andrew Murray, James Poulet and members of the Zampieri laboratory for comments on the manuscript. N.Z. and S.P. were supported by the DFG (ZA 885/1-1, ZA885/2-1 and EXC 257 NeuroCure).

## Material and Methods

### Mouse strains

Animals were housed in the facility with controlled environmental parameters under a 12h light/ 12h dark cycle and fed with standard chow. The following strains of mice were used in this study: *PV::Cre* (Hippenmeyer et al., 2007), *TRPV1::Cre* (Mishra et al.,2011), *TRPM8::Cre* (Yarmolinsky et al., 2006), *Rosa-lsl-HTB* (Li et al.,2013) and *Rosa-lsl-tdTomato* (Ai14, Jackson Laboratory). All animal experiments were approved by the Regional Office of Health and Social Affair Berlin (LAGeSo) and performed in compliance with the German Animal Walfare Act.

### Production of pseudotyped glycoprotein deficient rabies virus

RVΔG-mCherry/EnvA was produced with minor modifications as previously described (Wickersham et al., 2010). BHK-EnvA cells were infected with RVΔG-mCherry at a multiplicity of infection (MOI) of 2. 24 hours later cells were washed 3 times in PBS and fresh media added, this was repeated 24 hours later. After 48 hours incubation, media was harvested, filtered and viral particles concentrated by centrifugation. The virus was resuspended in PBS and further concentrated with Amicon Ultra 100 kDa protein concentrators. Viral titres were assessed by serial dilution of the virus on 293-TVA cells and virus of titre 1 × 10^8^ I.U./ml used for injection.

### Spinal cord injection

For rabies tracing experiments p9 *PV^HTB^*; *TRPV1^HTB^* and *TRPM8^HTB^* mice were anesthetized with isoflurane and placed on a stereotaxic frame. A skin incision in the back was made to expose the most caudal ribs to identify the lumbar spinal cord level 1. RVΔG-mCherry/EnvA was injected starting from 300 μm deep into the dorsal horn and going back dorsally, in 6 steps consisting of 50 nl pulses every 50 μm on the left side (400 μm lateral from the midline) of the spinal cord using a 0.5 μl Hamilton syringe mounted on a UMP3 UltraMicroPump (WPI). Skin was then sutured with a nylon surgical suture. Animals were sacrificed 7 days after injection (p16).

### Perfusion

Animals were anesthetized by intraperitoneal injection of 0.1 ml ketamine /xylazine mix per 10 g of weight (final concentrations: 120 mg/kg and 10 mg/kg, respectively) and checked for toe-pinch reflex before starting any procedure. Animals were first transcardially perfused with ice-cold PBS until the liver was cleared of blood, followed by ice-cold 4% PFA.

### Spinal cord dissection and tissue processing

Spinal cords were exposed by ventral laminectomy. Tissue was post-fixed overnight in 4% PFA at 4°C. This was followed by three washes with ice-cold PBS for 5 minutes each and over night incubation in 30% sucrose in phosphate buffer (0.1M PB) at 4°C for cryoprotection. Samples were embedded in Optimal Cutting Temperature (O.C.T., TissueTek) compound, frozen on dry ice and stored at −80°C.

### Immunohistochemistry

Consecutive sections (30μm thick) were made with a Leica cryostat and mounted on Superfrost Plus slides (VWR). For immunohistochemical staining, sections were hydrated with 1X PBS for 20 minutes and permeabilized with 0.1% Triton X-100/PBS for 10 minutes at room temperature. Primary antibodies diluted in Triton X-100/PBS were incubated overnight at 4°C. Primary antibody dilutions were used as follows: rabbit anti-DsRed 1:1000 (Takara), goat anti-ChAT 1:250 (Surmeli et al., 2011), sheep anti-GFP 1:2000 (Bio-rad), chicken anti-PV 1:10000 (de Nooij et al., 2013), sheep anti-Chx10 1:500 (Abcam), rabbit anti-Calbindin 28k 1:2000 (SWANT), goat anti-FoxP2 1:250 (Santa Cruz), guinea pig anti-Lbx1 1:10000 (Muller et al., 2002), rabbit anti-Lhx1 1:10000 (Generated in the Jessell laboratory), rabbit anti-PKCγ 1:500 (Cell Signaling Technology), FITC conjugated-IB4 (Sigma) and rabbit anti-CGRP 1:2000 (Immunostar). After washing 3 times with Triton X-100/PBS, sections were incubated with secondary antibodies for 1hour at room temperature. Alexa Fluor 488- and Cy3-conjugated secondary antibodies were used at 1:1000, Cy5-conjugated secondary antibodies at 1:500. Sections were then washed twice with 0.1% Triton X-100/PBS for 5 minutes each and once with 1X PBS for 10 minutes. Slides were coverslipped using Vectashield mounting medium. Images were acquired using confocal microscope (Zeiss LSM 800).

### Tissue clearing

The dura, from a post-fixed spinal cord, was carefully and completely removed. The tissue was cleared as previously described (Susaki et al., 2015). Briefly, samples were incubated at 37°C in ½ Scale CUBIC 1 with water for 3-6 hours and then incubated with Scale CUBIC 1 overnight at 37°C. On the 2nd day, the Scale CUBIC 1 was changed with a fresh one and leaved for other 2 days. Then, samples were washed with 1X PBS overnight and incubated in ½ Scale CUBIC 2 in PBS for 3-6 hours at 37°C. The next day, samples were transferred in Pure Scale CUBIC 2 overnight at 37°C. After clearing, samples were immediately imaged in mineral oil with a Zeiss Z1 light sheet miscroscope.

### Neuronal position analysis

Three-dimensional positional analysis was performed as previously described (Dewitz et al., 2018). Briefly, high-resolution images of the spinal cord were processed with the imaging software IMARIS using the “spots” function to assign Cartesian coordinates to all labeled neurons. We set the central canal as the 0, 0 coordinate for the medio-lateral (x-axis) and dorso-ventral (y-axis) axes. These coordinates (x and y) were rotated and normalized to a standardized spinal cord, whose dimensions were obtained by calculating the average size of spinal cords at p16 (M-L: 800 um, D-V: 600 um), to avoid variability in size and orientation of the spinal cord between experiments. Datasets were aligned on the z-axis by starting analysis from the section where the first labeled neurons appeared (z=0) in the L1 segment and progressed caudally for more than 2 mm, covering two lumbar segments of the spinal cord. Positional analyses were performed using custom script in “R project” (R Foundation for Statistical Computing, Vienna, Austria, http://www.r-project.org). Contour and Density plot were generated using “ggplot2” package. The heat maps were used to compare the 2D spatial distribution of interneurons within each experiment and generated with the “corrplot” function. The similarity between experiments was measured by the Pearson correlation coefficient “r”.

