## Supplementary figures and images for "Rabies anterograde monosynaptic tracing reveals organization of spinal sensory circuits"

### Supplemental Figure 1

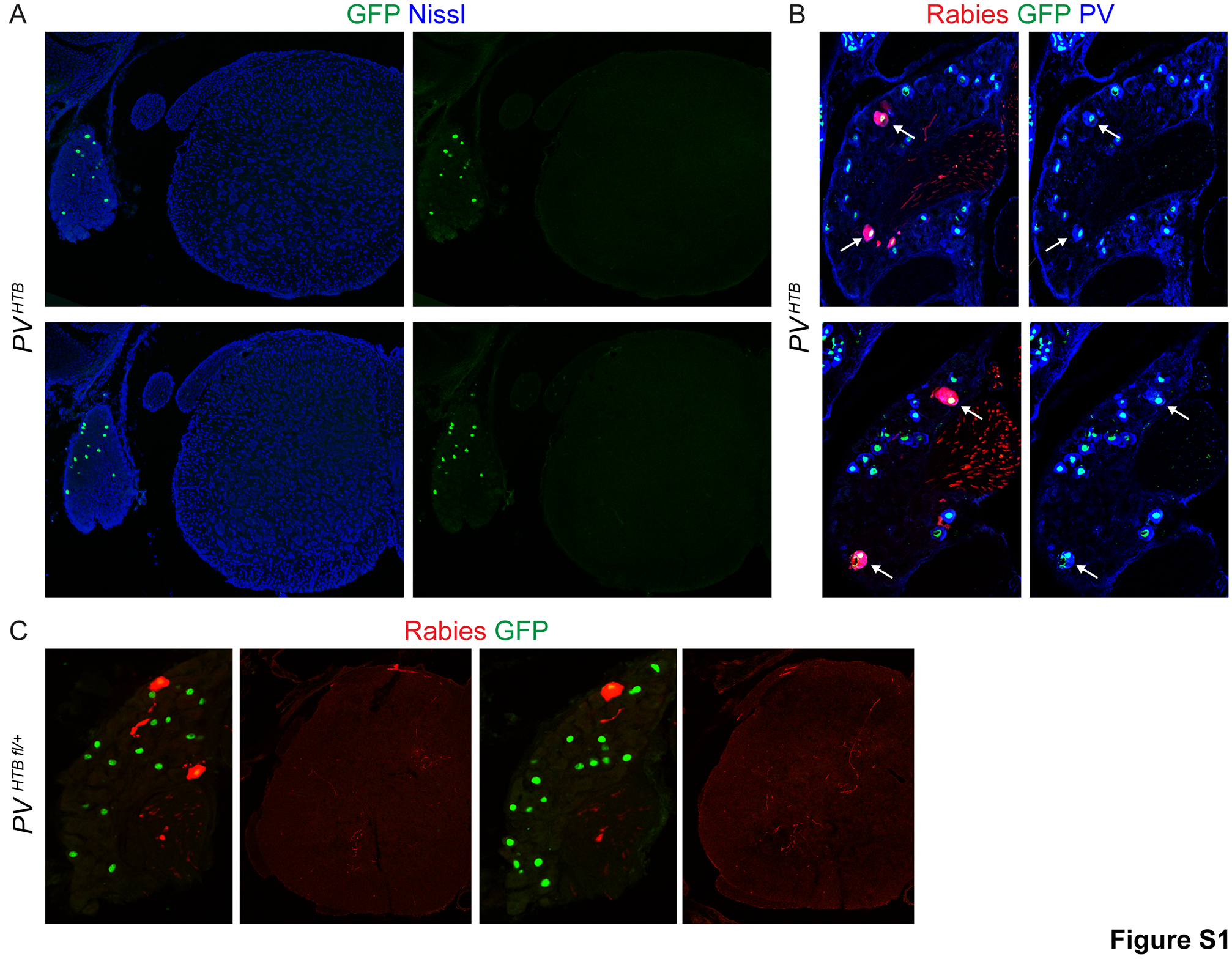

### Supplemental Figure 2

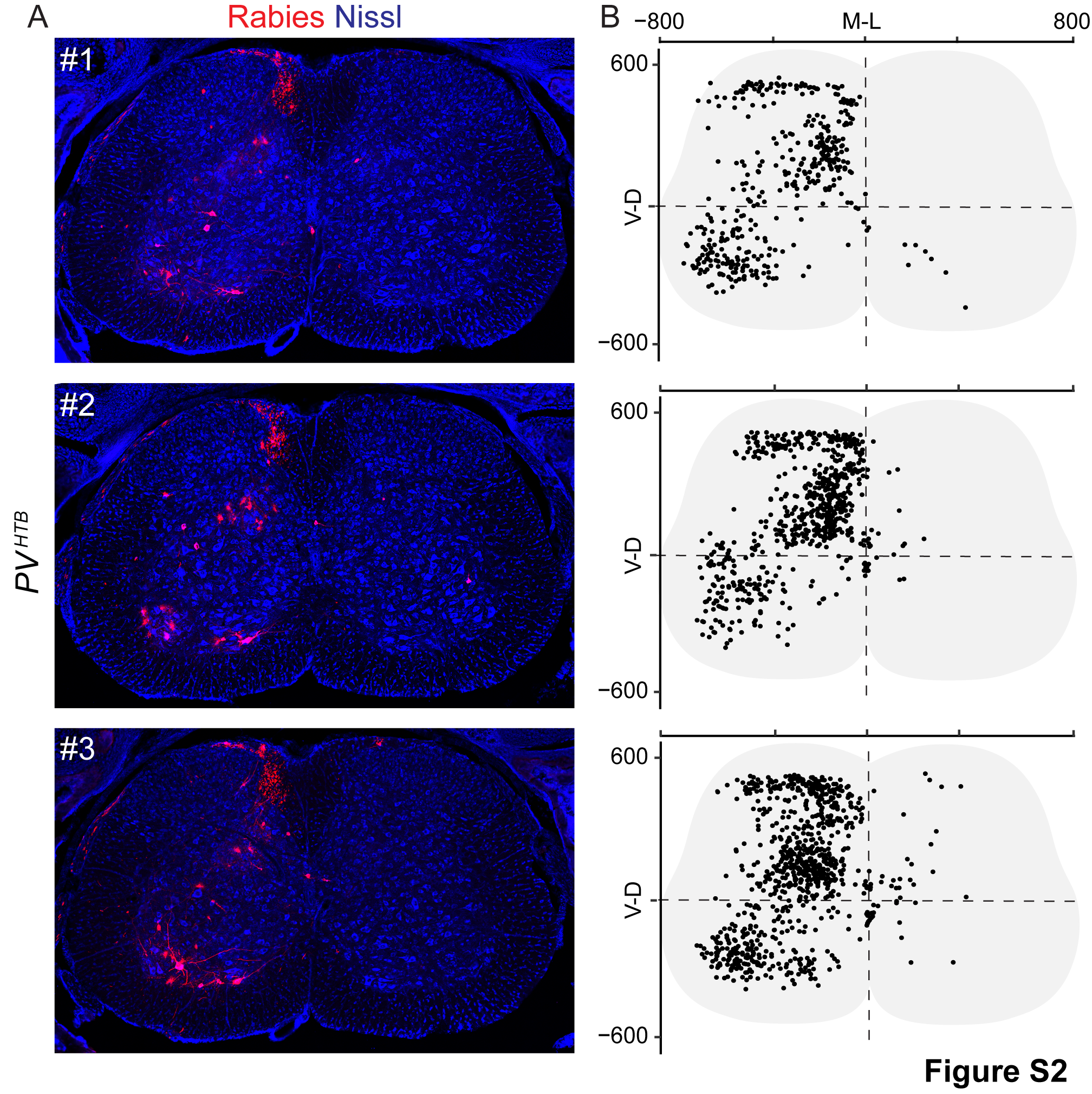

### Supplemental Figure 3

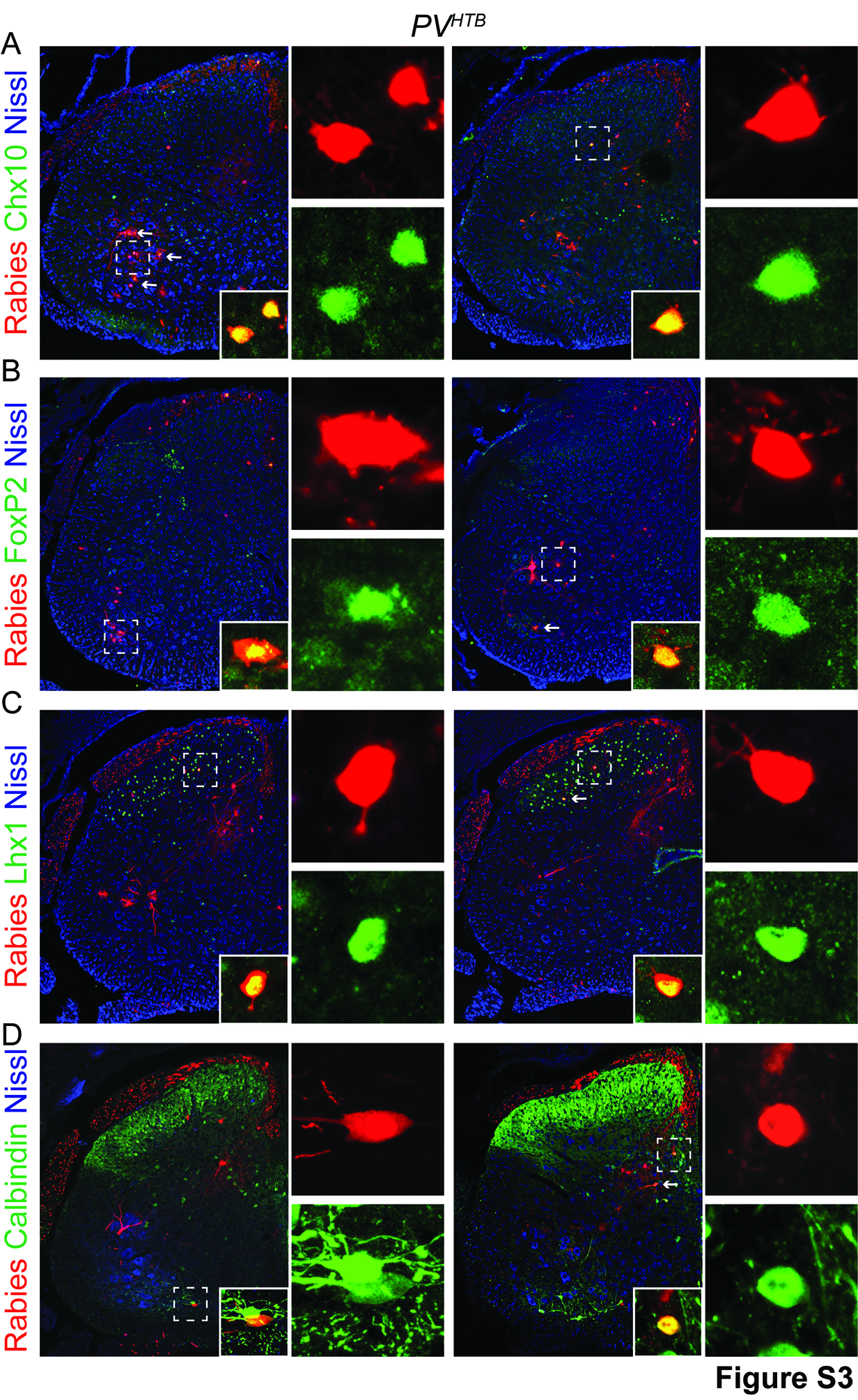

### Supplemental Figure 4

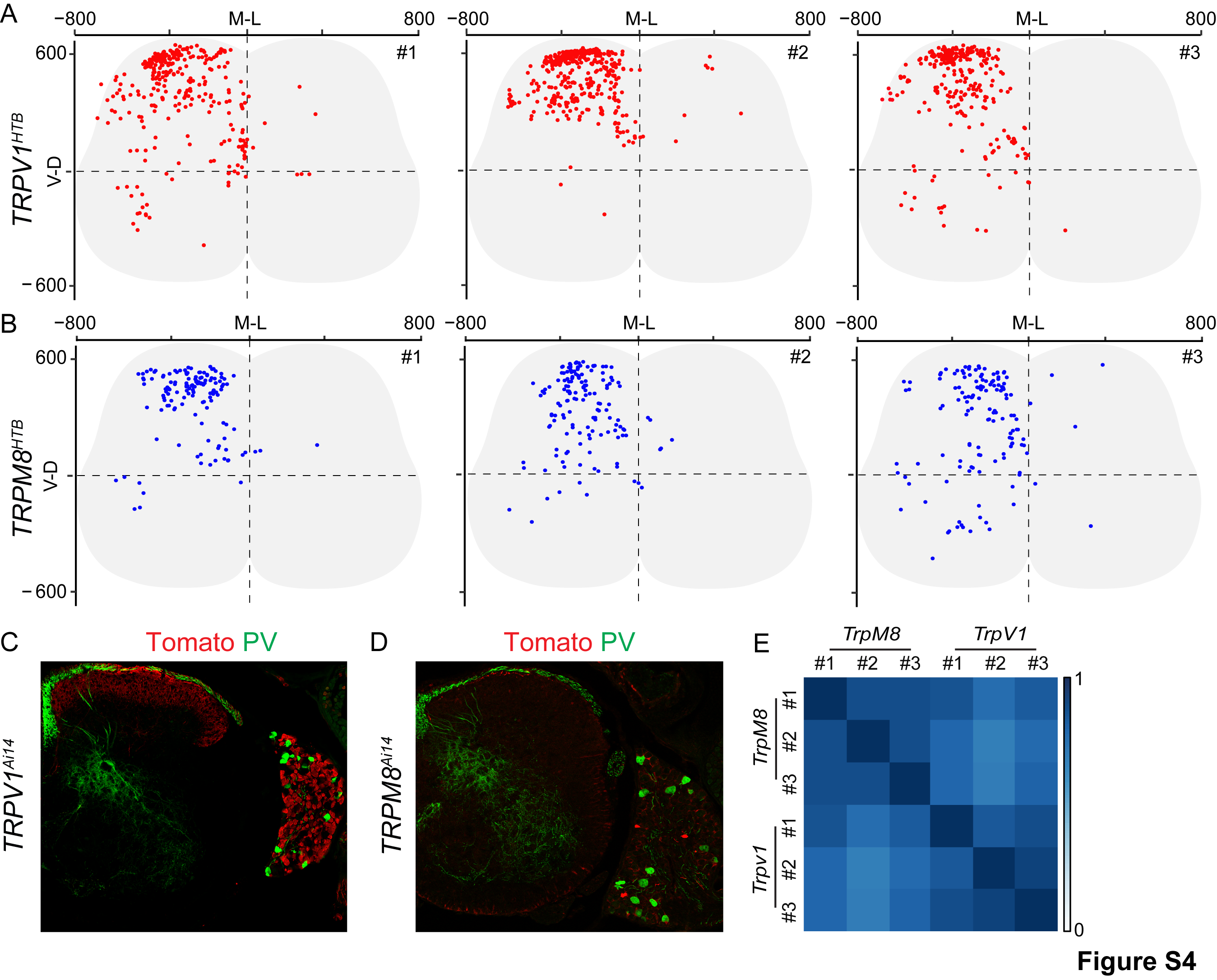
