## Supplemental Figure Legends for "Rabies anterograde monosynaptic tracing reveals organization of spinal sensory circuits"

**Figure S1. Specific infection of proprioceptive neurons with *PV^HTB^* mouse line.**

A) Representative image of a lumbar spinal cord section showing GFP^+^ sensory neurons in the DRG of p10 *PV^HTB^* mice.

B) Examples of RV^+^; GFP^+^; PV^+^ sensory neurons after RV∆G-mCherry/EnvA L1 injection in p9 *PV^HTB^* mice.

C) Representative images of lumbar DRG and spinal cord sections showing RV^+^; GFP^+^ sensory neurons and absence of transsynaptic labeling after RV∆G-mCherry/EnvA L1 injection in p9 *PV::cre^+/-^; HTB^f/+^* (*PV^HTB f/+^*) mice.

**Figure S2. Post-sensory connectivity maps from *PV^HTB^* experiments.**

A) RV^+^ spinal neurons after RV∆G-mCherry/EnvA L1 injection in three p9 *PV^HTB^* mice.

B) Digital reconstruction of RV^+^ neuron positions in three *PV^HTB^* experiments.

**Figure S3. Subtype identities of post-sensory neurons labeled in *PV^HTB^* experiments.**

A-D) Representative images of Chx10^+^ (V2a, A), FoxP2^+^ (V1, B), Lhx1^+^ (V0/dI4, C) and calbindin^+^ (D) interneurons labeled in *PV^HTB^* experiments.

**Figure S4. Post-sensory connectivity maps from *TRPV1^HTB^* and *TRPM8^HTB^* experiments.**

A and B) Digital reconstruction of RV^+^ neuron positions in three *TRPV1^HTB^* (A, red)) and *TRPM8^HTB^* (B, blue) experiments.

C and D) Representative images of parvalbumin and tdTomato labeling of somatosensory neurons cell bodies and afferents in the DRG and spinal cord of *TRPV1::cre^+/-^; Ai14^f/+^* (C) and *TRPM8::cre^+/-^; Ai14^f/+^* mice.

E) Correlation analysis of post-sensory neurons Cartesian coordinates in *TRPV1^HTB^* and *TRPM8^HTB^* experiments.
