## Supplemental Table 1 for "Rabies anterograde monosynaptic tracing reveals organization of spinal sensory circuits"

| **Experiment** | **Animal** | **#** | **# of starter cells** | **# of spinal neurons** | **Specificity (%)** | **Efficiency (%)** | **Connectivity index** |
| --- | --- | --- | --- | --- | --- | --- | --- |
|  | *PV:: cre +/-; Rosa-HTB f/f* | *1* | 85 | 413 | 89.0 | 12 | 4.86 |
| *PV^HTB^* | *PV:: cre +/-; Rosa-HTB f/f* | *2* | 141 | 696 | 94.0 | 14 | 4.94 |
|  | *PV:: cre +/-; Rosa-HTB f/f* | *3* | 194 | 858 | 94.0 | 16 | 4.42 |
|  | *TRPV1::cre+/-; Rosa-HTB f/f* | *1* | 2939 | 545 | 98.5 | 17 | 0.19 |
| *TRPV1^HTB^* | *TRPV1::cre+/-; Rosa-HTB f/f* | *2* | 1580 | 302 | 98.0 | 30 | 0.19 |
|  | *TRPV1::cre+/-; Rosa-HTB f/f* | *3* | 1642 | 312 | 98.2 | 13 | 0.19 |
|  | *TRPM8::cre+/-; Rosa-HTB f/f* | *1* | 7 | 139 | 75.0 | 40 | 19.86 |
| *TRPM8^HTB^* | *TRPM8::cre+/-; Rosa-HTB f/f* | *2* | 8 | 154 | 85.0 | 38 | 19.25 |
|  | *TRPM8::cre+/-; Rosa-HTB f/f* | *3* | 7 | 143 | 80.0 | 32 | 20.42 |
